## Supplementary material for "Synthetic Aptamer Mechanoreceptors Enable Cell-Specific Force Sensing and Temporal Control via DNA Circuits": SI

**Supplementary Tables**

**Supplementary Table 1.** DNA sequences for the oligonucleotides with their names, sequence codes, and modifications.

| Name | Sequence 5'-3' | Modification |
|---|---|---|
| Unzipping mode | AGT CGT ATT ACC GCG TTT | 5' Rhodamin Red - X<br>3' Biotin-TEG |
| Shearing mode | TTT 8AG TCG TAT TA CCGCG | 5' Biotin-TEG<br>8 dT-Rhodamine Red-X |
| AS1411 strand | GGT GGT GGT GGT TGT GGT GGT GGT GGC GCG GTA ATA CGA CT | 3' BHQ2 |
| AS1411 strand with Atto 488 | TTG GTG GTG GTG GTT GTG GTG GTG GTG GCG CGG TAA TAC GAC T | 5' Atto 488 |
| Random strand | CCT CCT CCT CCT TCT CCT CCT CCT CCC GCG GTA ATA CGA CT | 3' BHQ2 |
| Null strand | CGC GGT AAT ACG ACT | 3' BHQ2 |
| TB-5 strand | TTT TTA AGC CCA CTC CTC TGT GGG GGG CGA ACA ACA AGG CAG TCG TGC CAT GCG CGG TAA TAC GAC T | 3' BMNQ1 |
| DBCO strand [a] | TTC GCG GTA ATA CGA CT | 5' DBCO<br>3' BHQ2 |
| Sgc8 strand | ATC TAA CTG CTG CGC CGC CGG GAA AAT ACT GTA CGG TTA GAC GCG GTA ATA CGA CT | 3' BHQ2 |
| MUC1 strand | GCA GTT GAT CCT TTG GAT ACC CTG GCG CGG TAA TAC GAC T | 3' BHQ2 |
| SYL3c strand | CAC TAC AGA GGT TGC GTC TGT CCC ACG TTG TCA TGG GGG GTT GGC CTG CGC GGT AAT ACG ACT | 3' BHQ2 |
| DNA Blocker strand | TAC GGC GAG ACA CCA CCA CCA CCA CAA CCA CCA CCA CC | 3' BHQ1 |
| Activator strand | GGT GGT GGT GGT TGT GGT GGT GGT GGT GTC TCG CCG TA | |
| RNA blocker strand | CCA CCA CCA CCA CAA CCA CCA CCA CC | 3' BHQ1 |
| Random RNA strand | GCA UUU CGU CUU AAU AUA GCA UGU CA | |
| Random DNA strand | TGA CAT GCT ATA TTA AGA CGA AAT GC | |
| AS1411 strand without quencher | GGT GGT GGT GGT TGT GGT GGT GGT GGC GCG GTA ATA CGA CT | |
| MUC1 strand without quencher | GCA GTT GAT CCT TTG GAT ACC CTG GCG CGG TAA TAC GAC T | |
| Sgc8 strand without quencher | ATC TAA CTG CTG CGC CGC CGG GAA AAT ACT GTA CGG TTA GAC GCG GTA ATA CGA CT | |
| SYL3c strand without quencher | CAC TAC AGA GGT TGC GTC TGT CCC ACG TTG TCA TGG GGG GTT GGC CTG CGC GGT AAT ACG ACT | |
| DBCO strand without quencher | TT CGCGG TAA TAC GAC T | 5' DBCO |

[a] We used click chemistry to conjugate cRGDfK-$N_3$ to DBCO strand for obtaining RGD strand based on our reported method.[1]

**Supplementary Table 2.** Surface density of aptamer MPs and RGD molecules. n = 18 from 3 replicates.

| | **MP density (molecule/μm$^2$)** | **RGD-bio density (molecule/μm$^2$)[a]** |
|---|---|---|
| AS1411 unzipping | 4329±394 | 1297±155 |
| Sgc8 unzipping | 4480±239 | 1320±159 |
| MUC1 S2.2 unzipping | 4040±216 | 1322±101 |
| SYL3c unzipping | 4072±166 | 1356±134 |
| AS1411 shearing | 4141±269 | 1254±82 |
| Sgc8 shearing | 3918±609 | 1321±61 |
| RGD unzipping | 3495±201 | 1374±92 |
| RGD shearing | 3581±241 | 1318±155 |

**Supplementary Table 3.** Reaction rate constant of SDR and RNA-RNase H modules. To understand the reconfiguration efficiency and tunability of different DNRs, we quantified the surface reconfiguration kinetics. The data were analyzed assuming pseudo-first-order kinetics and fitted using one phase exponential decay function. For the SDR module, reactivation with 200 nM DNA activator yields a reconfiguration rate of $2.14 \times 10^{-2}$ $min^{-1}$. The RNA-RNase H module enables degradation rate tuning by adjusting the initial RNase H concentration, resulting in accelerated (100 U/mL, $5.26 \times 10^{-2}$ $min^{-1}$) or decelerated (10 U/mL, $0.71 \times 10^{-2}$ $min^{-1}$) reconfiguration kinetics. Moreover, at a fixed RNase H concentration (10 U/mL), the RNA-RNase H module allows the introduction of non-enzymatic RNA/DNA decoys, providing an additional orthogonal layer of regulation.

| | ***k* ($\times 10^{-2}$ $min^{-1}$)** | **$R^2$** |
|---|---|---|
| SDR 200 nM DNA activator | 2.14 | 0.9926 |
| RNase H 100U/mL | 5.26 | 0.9925 |
| RNase H 10U/mL | 0.71 | 0.9842 |
| RNase H 10U/mL+1 nM rdm RNA/DNA decoy | 0.73 | 0.9782 |
| RNase H 10U/mL+10 nM rdm RNA/DNA decoy | 0.42 | 0.9603 |
| RNase H 10U/mL+100 nM rdm RNA/DNA decoy | 0.11 | 0.9831 |

### Supplementary Figures

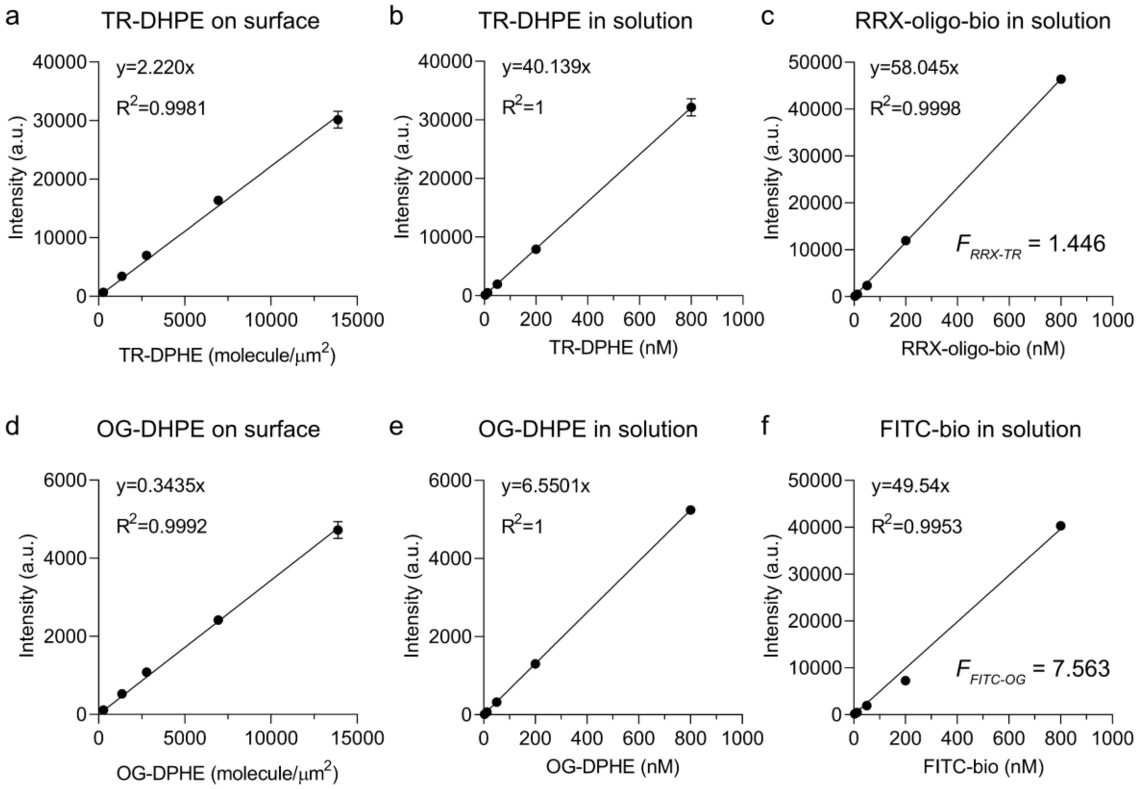


**Supplementary Figure 1.** Quantification of surface density of aptamer MPs and RGD molecules on the surface. **a-c** Standard curves for quantifying density of aptamer MPs. **a** SLB calibration: Intensity of Texas Red DHPE (Texas Red™ 1,2-dihexadecanoyl-sn-glycero-3-phosphoethanolamine, triethylammonium salt) to the number of molecules on the surface. n = 15 from 3 replicates. **b-c** F-factor plot estimation using concentrations of TR-DHPE-derived SUVs (**b**) and RRX-labelled oligonucleotide in solution (**c**) to compare the fluorescence intensity with density. The ratio of the calibration curve slopes was used to determine the "*F* factor" for the labelled oligonucleotide and the SUV samples. n = 15 from 3 replicates. **d-f** Standard curves for quantifying density of RGD molecules. Direct quantification of the pristine c[RGDfK(Biotin)] molecule is unfeasible due to its lack of a fluorophore. Using a fluorophore-conjugated RGD-biotin variant may distort the actual RGD density due to the increase in size and steric hindrance. Thus, we chose FITC-biotin (Fluorescein-5(6)-biotinamidohexanoylamidopentylthiourea, Mw 831.01 g/mol) as a proxy to estimate the RGD density, given its similar molecular weight to c[RGDfK(Biotin)] (Mw 829.98 g/mol). It should be noted that this approach ignores potential effects arising from differences in charge and hydrophilicity/hydrophobicity. **d** SLB calibration: Intensity of Oregon Green 488 DHPE (Oregon Green™ 488 1,2-Dihexadecanoyl-*sn*-Glycero-3-Phosphoethanolamine) to the number of molecules on the surface. n = 12 from 3 replicates. **e-f** F-factor plot estimation using concentrations of OG-DHPE-derived SUVs (**e**) and FITC-biotin in solution (**f**) to compare the fluorescence intensity with density. The ratio of the calibration curve slopes was used to determine the "*F* factor" for the labelled oligonucleotide and the SUV samples. n = 12 from 3 replicates.

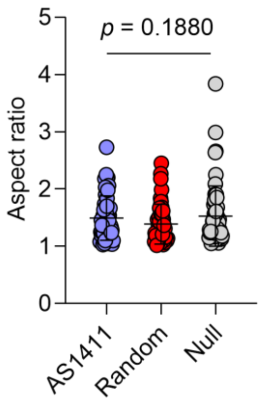


**Supplementary Figure 2.** Quantification of aspect ratio of HeLa cells on AS1411 MP surfaces. n = 54 cells from 3 replicates. Statistics: Kruskal-Wallis test with Dunn's multiple comparisons.

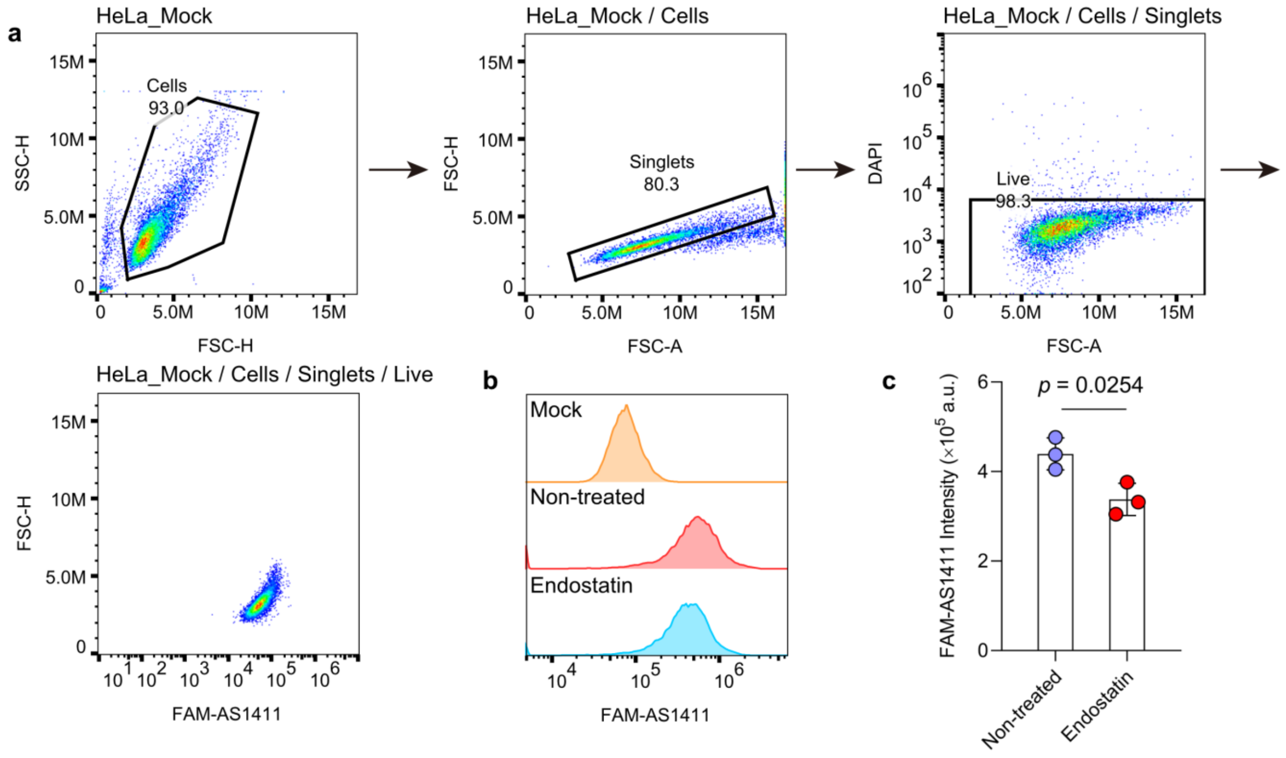


**Supplementary Figure 3. a** Flow cytometry gating strategy. **b** Representative flow cytometry histogram showing HeLa cells incubated with 2.5 µM FAM-AS1411 at 4 °C for 60 min, either in the absence or presence of 25 µg/mL endostatin. The mock control represents cells without FAM-AS1411, indicating cellular autofluorescence. **c** Median fluorescence intensity of FAM-AS1411 in HeLa cells after endostatin treatment. Endostatin significantly inhibits AS1411 binding to HeLa cells. n = 3 independent experiments. Statistics: unpaired two-tailed Student's *t*-test.

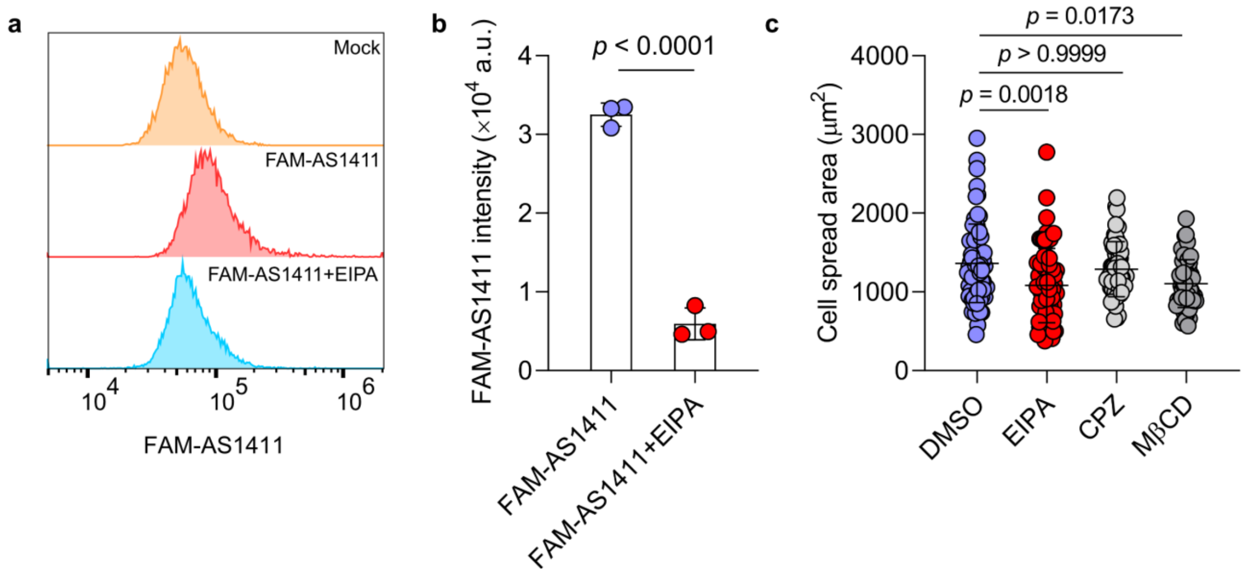


**Supplementary Figure 4. a** Representative flow cytometry histogram showing uptake of 500 nM FAM-AS1411 by HeLa cells at 37 °C for 4 h, either in the absence or presence of 100 μM EIPA. The mock control represents cells without FAM-AS1411, indicating cellular autofluorescence. **b** Median fluorescence intensity of FAM-AS1411 in HeLa cells after EIPA treatment. EIPA significantly inhibits AS1411 internalization by HeLa cells. n = 3 independent experiments. Statistics: unpaired two-tailed Student's *t*-test. **c** Quantification of cell spreading area when HeLa cells were treated with different internalization inhibitors on AS1411 MP surfaces. n = 70, 70, 54, 54 cells from 3 replicates. Statistics: Kruskal-Wallis test with Dunn's multiple comparisons.

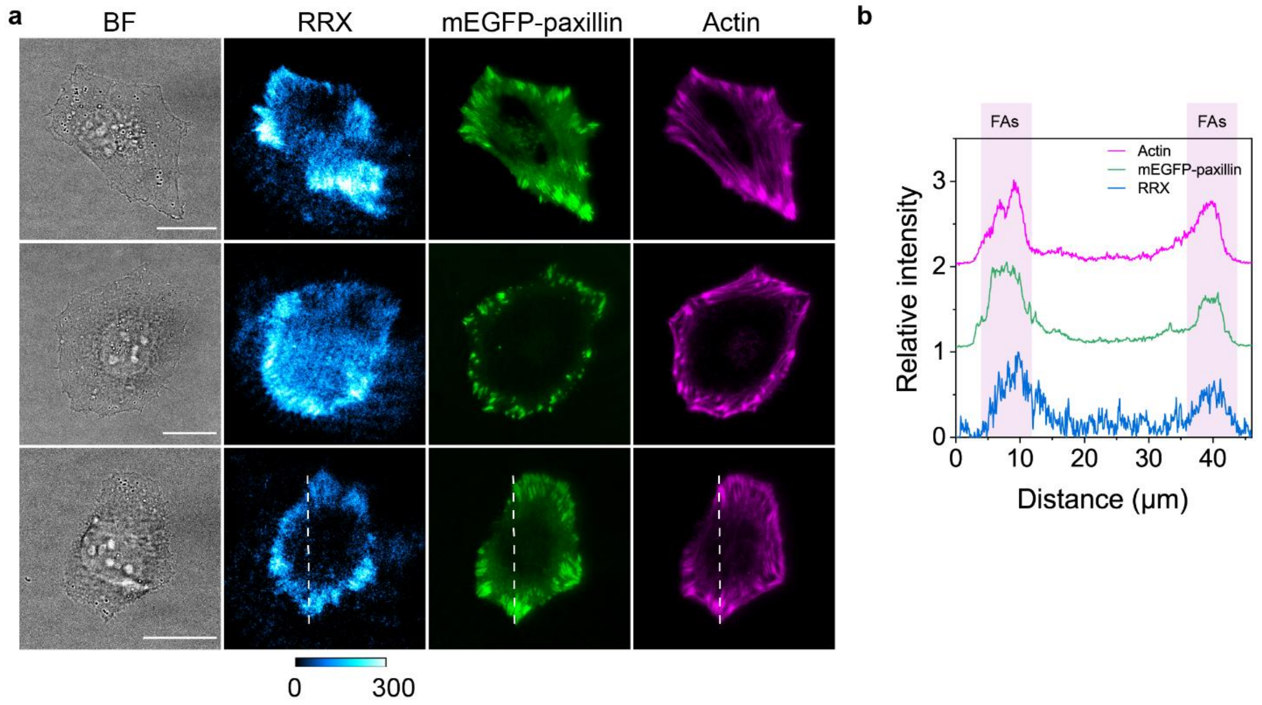


**Supplementary Figure 5. a** Representative microscopy imaging of brightfield, mechanosiganls, mEGFP-paxillin, and actin of HeLa cells incubated on AS1411 MP surfaces. **b** Line profile shows the intensity profiles of mEGFP-paxillin, actin and mechanosiganls along the white line marked in **a**. Nucleolin-generated forces are clearly localized at peripheral FA sites marked by paxillin. Scale bars = 20 μm.

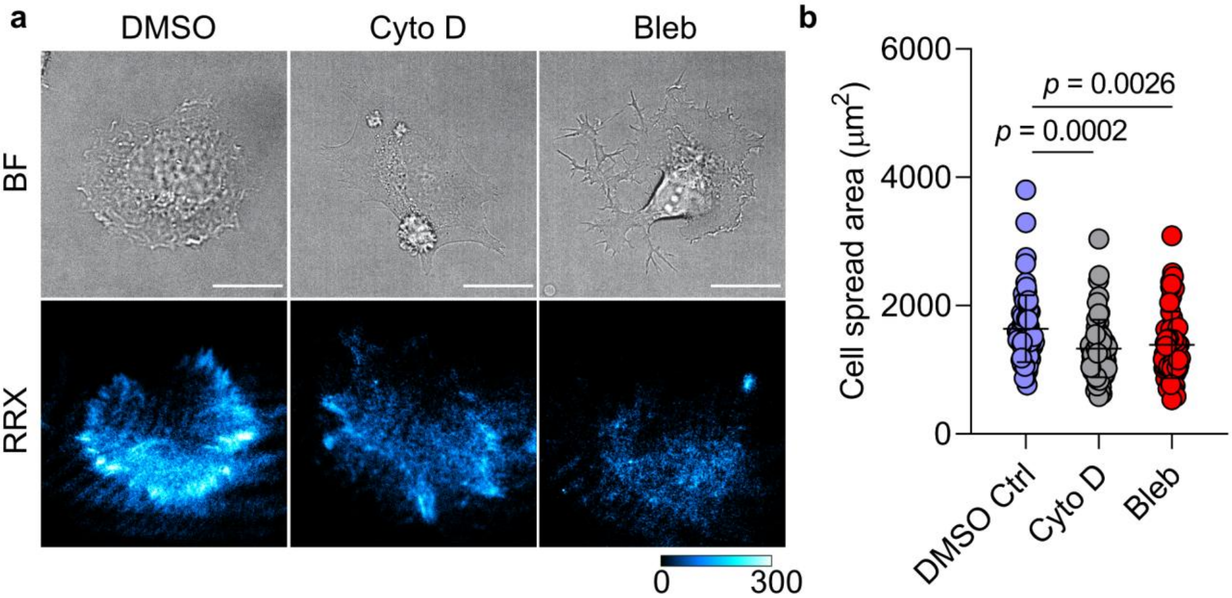


**Supplementary Figure 6. a** Representative brightfield and fluorescence images of HeLa cells treated with different cytoskeleton inhibitors on AS1411 MP surfaces. **b** Quantification of cell spreading area when HeLa cells were treated with different cytoskeleton inhibitors on AS1411 MP surfaces. Both cytochalasin D and blebbistatin reduce cell spreading area. n = 72 cells from 3 replicates. Statistics: Kruskal-Wallis test with Dunn's multiple comparisons. Scale bars = 20 µm.

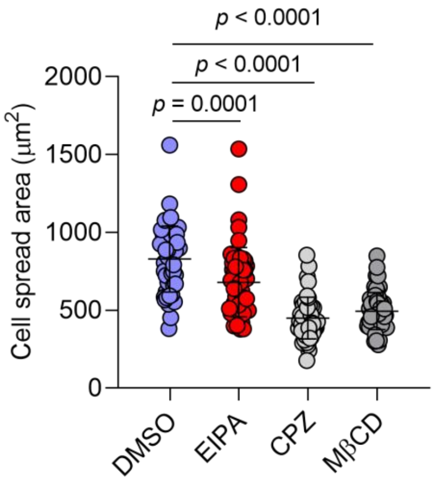


**Supplementary Figure 7.** Quantification of cell spreading area when HepG2 cells were treated with different internalization inhibitors on Sgc8 MP surfaces. n = 54 cells from 3 replicates. Statistics: One-way ANOVA with Bonferroni post-hoc tests.

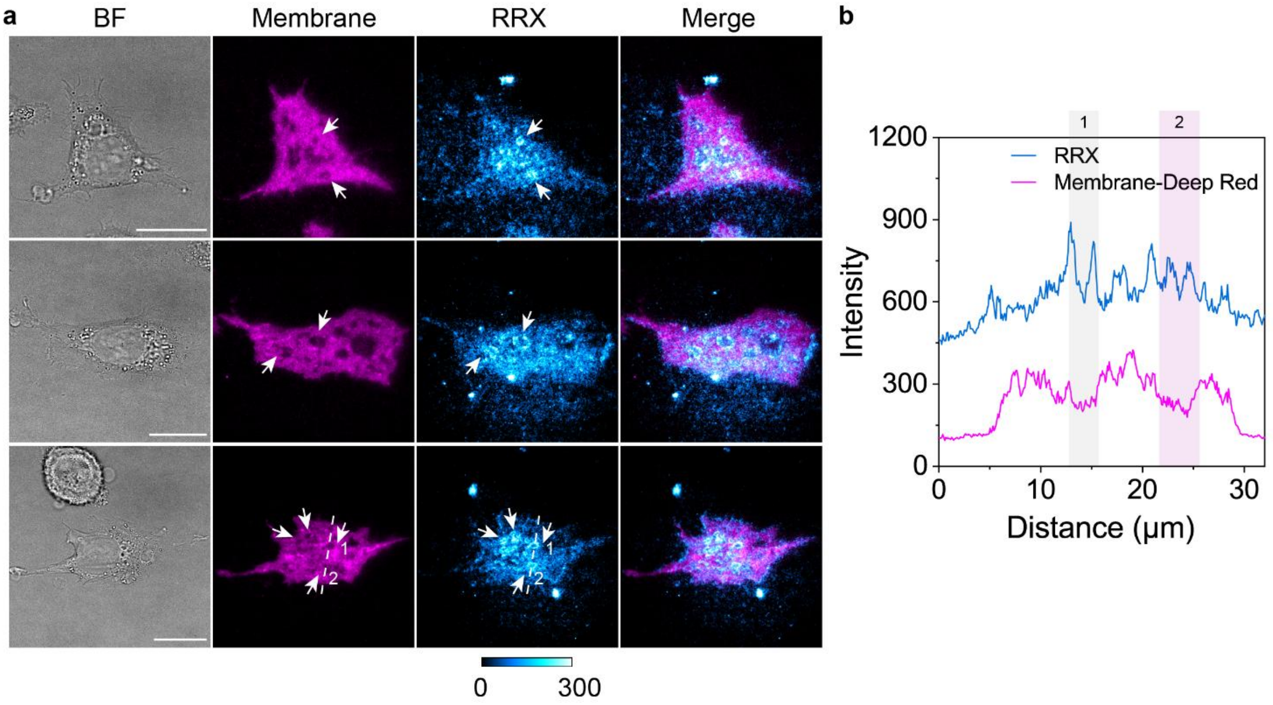


**Supplementary Figure 8. a** Representative microscopy imaging of brightfield, plasma membrane, and mechanosignals of HepG2 cells incubated on Sgc8 MP surfaces. **b** Line profile shows the intensity profiles of plasma membrane and mechanosignals along the white line marked in **b**. Mechanosignals are localized at the inner edge of membrane invaginations. Scale bars = 20 μm.

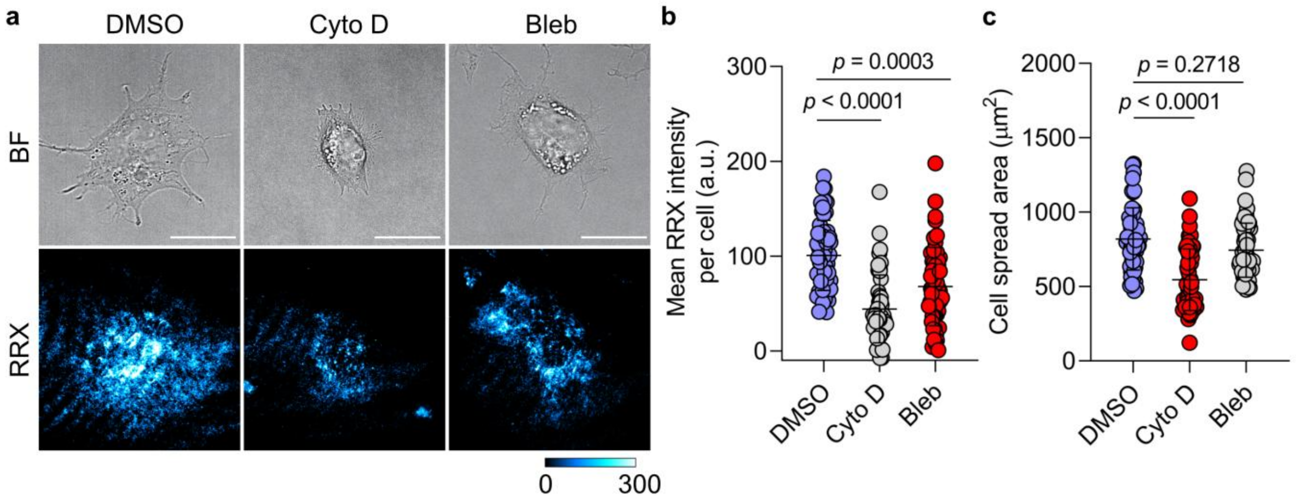


**Supplementary Figure 9. a** Representative brightfield and fluorescent images of HepG2 cells treated with different cytoskeleton inhibitors on Sgc8 MP surfaces. **b-c** Quantification of mean fluorescence intensity per cell (**b**) and cell spreading area (**c**) when HepG2 cells were treated with different cytoskeleton inhibitors on Sgc8 MP surfaces. Cytochalasin D (actin polymerization inhibitor) exerts a stronger inhibitory effect than blebbistatin (myosin II inhibitor), indicating that actin polymerization plays a more improtant role than actomyosin contractility in PTK7-mediated force generation. n = 54 cells from 3 replicates. Statistics: Kruskal-Wallis test with Dunn's multiple comparisons. Scale bars = 20 μm.

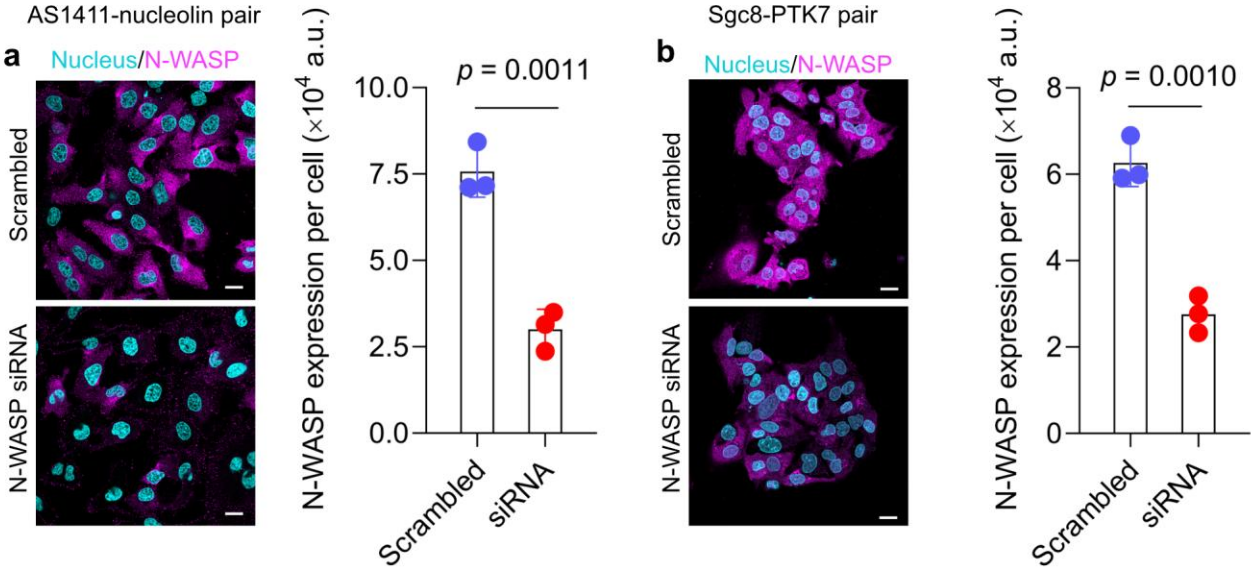


**Supplementary Figure 10. a** Representative immunostaining images and quantification of N-WASP expression level of HeLa cells after siRNA knockdown. HeLa cells transfected with either scrambled or N-WASP siRNA were stained with anti-N-WASP primary antibody followed by Alexa Fluor™ 647-conjugated secondary antibody. n = 1372 and 1304 cells from 3 replicates for scrambled and siRNA groups. Statistics: unpaired two-tailed Student's *t*-test. **b** Representative immunostaining images and quantification of N-WASP expression level of HepG2 cells after siRNA knockdown. HepG2 cells transfected with either scrambled or N-WASP siRNA were stained with anti-N-WASP primary antibody followed by Alexa Fluor™ 647-conjugated secondary antibody. n = 1222 and 1198 cells from 3 replicates for scrambled and siRNA groups. Statistics: unpaired two-tailed Student's *t*-test. Scale bars = 20 µm.

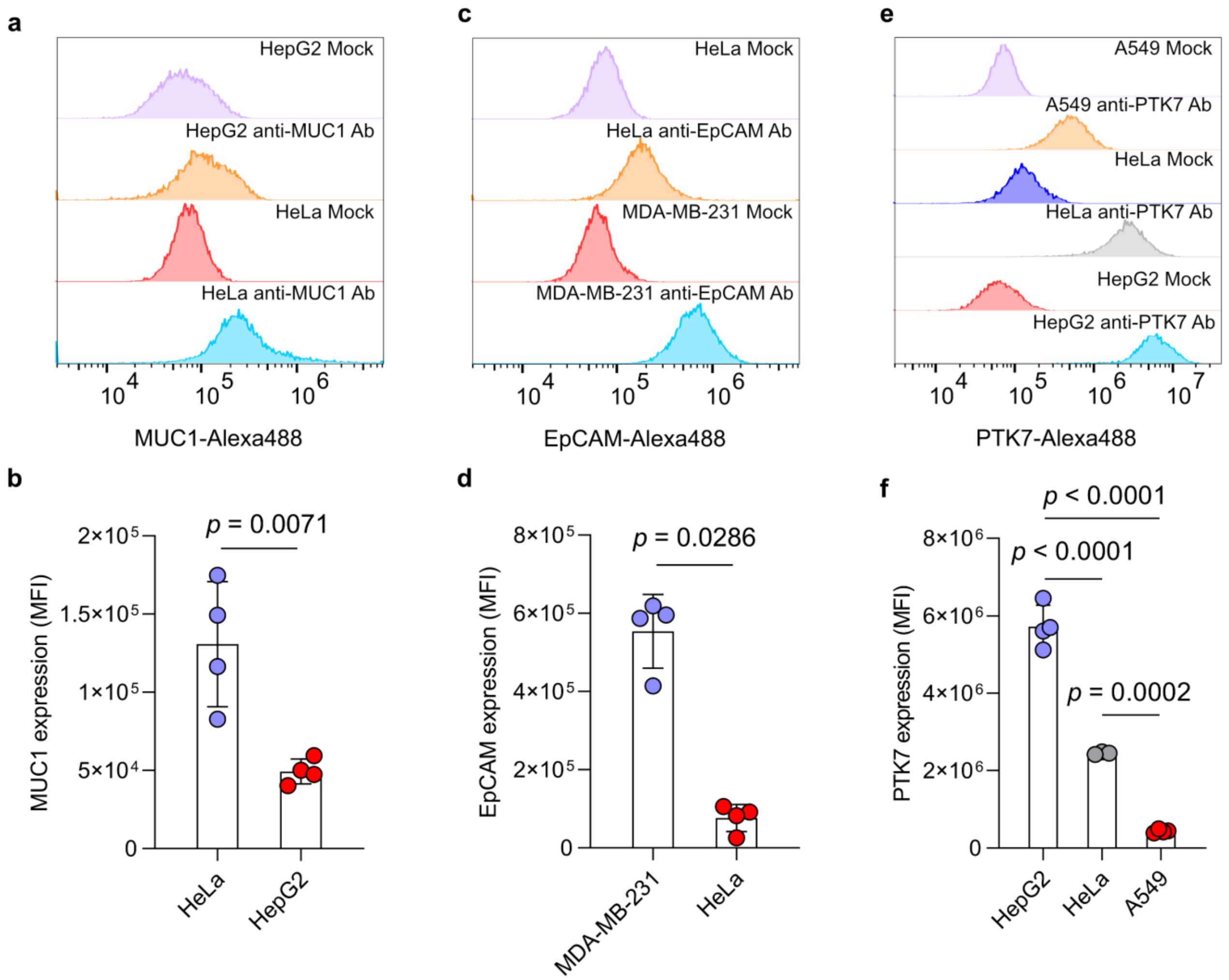


**Supplementary Figure 11.** **a** Representative flow cytometry histogram of HeLa and HepG2 cells stained with Alexa Fluor™ 488-conjugated anti-MUC1 primary. **b** Quantification of mucin-1 expression levels on HeLa and HepG2 cells based on median fluorescence intensity. n = 3 independent experiments. Statistics: unpaired two-tailed Student's *t*-test. **c** Representative flow cytometry histogram of MDA-MB-231 and HeLa cells stained with Alexa Fluor™ 488-conjugated anti-EpCAM primary. **d** Quantification of EpCAM expression levels on MDA-MB-231 and HeLa cells based on median fluorescence intensity. n = 3 independent experiments. Statistics: two-tailed, Mann-Whitney test. **e** Representative flow cytometry histogram of HepG2, HeLa, and A549 cells stained with anti-PTK7 primary antibody followed by Alexa Fluor™ 488-conjugated secondary antibody. Mock controls represent unstained cells, indicating cellular autofluorescence. **f** Quantification of PTK7 expression levels on HepG2, HeLa, and A549 cells based on median fluorescence intensity. n = 3 independent experiments. Statistics: One-way ANOVA with Bonferroni post-hoc tests.

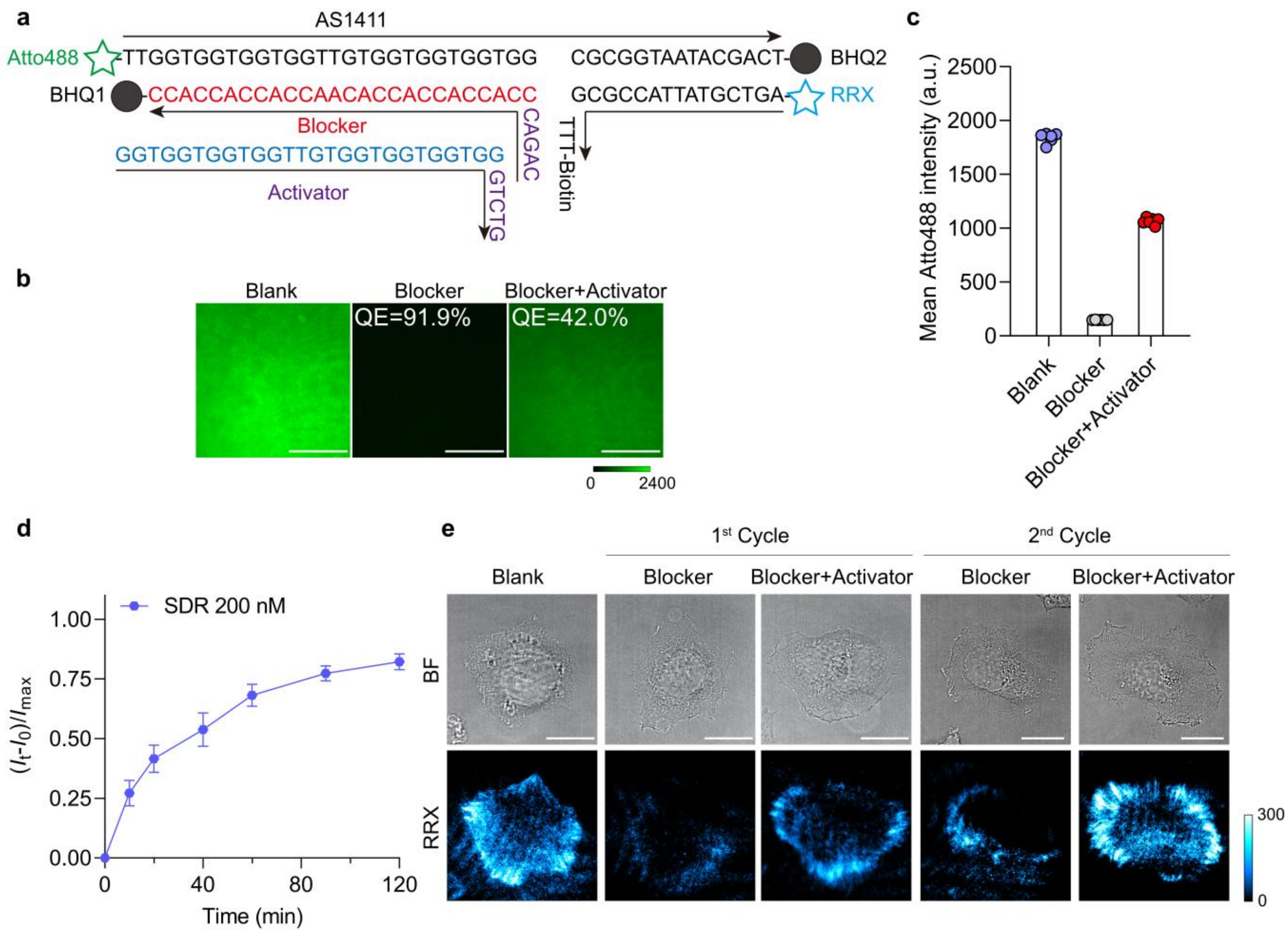


**Supplementary Figure 12. a** Detailed sequences for controlling AS1411 MPs by SDRs. **b-c** Fluorescence images (**b**) and corresponding quantification (**c**) showing blocking and reactivation of the surface-immobilized AS1411 MP. MPs are labeled with Atto488 at the 5' end. Incubation with 200 nM 3'-BHQ1–labeled blocker strands for 30 min results in a 91.9% decrease in Atto488 fluorescence. Subsequent medium exchange with the equimolar activator restores fluorescence by 49.9% within 30 min. Reaction medium: 1 % FBS, 1 % P/S, DMEM. **d** Plot of normalized surface Atto488 intensity versus time to compare the reconfiguration rate of AS1411 aptamer in SDR module. Surface-immobilized AS1411 MPs are initially blocked with DNA blocker and subsequently reconfigured by adding 200 nM DNA activator. At each reaction time point, the surface Atto488 intensity is normalized to that of unblocked AS1411 MPs labeled with Atto488, which serves as the positive control. Reaction medium: 1 % FBS, 1 % P/S, 100 µg/mL actin protein, DMEM. n = 15 from 3 replicates. **e** Representative brightfield and fluorescence images of HeLa cells on AS1411 MP surfaces after sequential blocker and activator treatment. Scale bars = 20 µm.

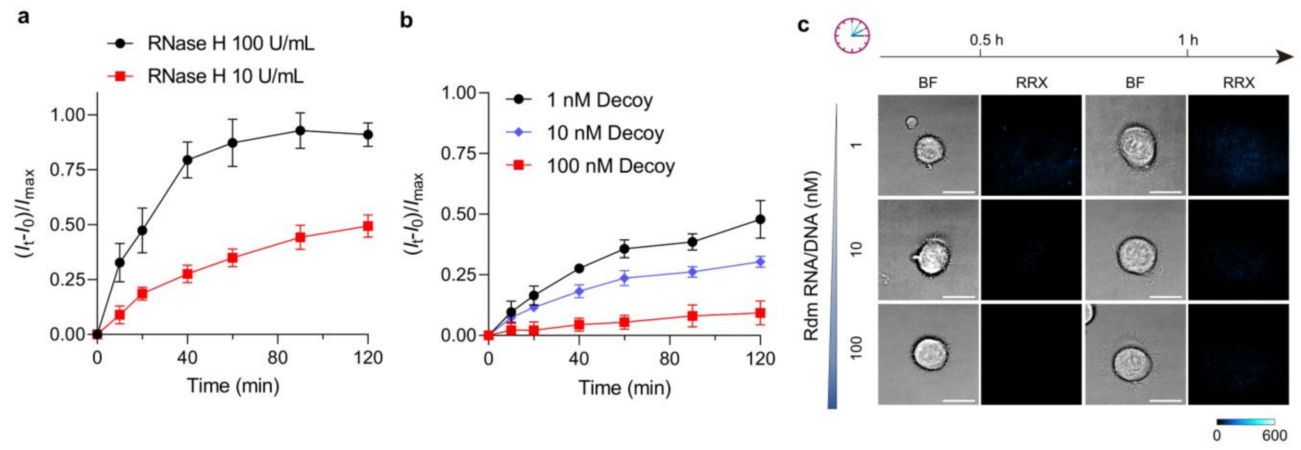


**Supplementary Figure 13. a** Plot of normalized surface Atto488 intensity versus time to compare the reconfiguration rate of AS1411 aptamer in RNA-RNase H module. Surface-immobilized AS1411 MPs are blocked with RNA blocker and degraded by RNase H at different concentrations (10 or 100 U/mL). At each reaction time point, the surface Atto488 intensity is normalized to that of unblocked AS1411 MPs labeled with atto488, which serves as the positive control. Reaction medium: 1 % FBS, 1 % P/S, 100 µg/mL actin protein, DMEM. n = 15 from 3 replicates. **b** Plot of normalized surface Atto488 intensity versus time to compare the reconfiguration rate of AS1411 aptamer in RNA-RNase H module in the presence of decoy strands. Surface-immobilized AS1411 MPs are blocked with RNA blocker and degraded by RNase H at 10 or 100 U/mL and decoy strands at 1-100 nM. Reaction medium: 1 % FBS, 1 % P/S, 100 µg/mL actin protein, DMEM. n = 15 from 3 replicates. At each reaction time point, the surface Atto488 intensity is normalized to that of unblocked AS1411 MPs labeled with atto488, which serves as the positive control. **c** Representative brightfield and fluorescence images of HeLa cells on RNA-blocked AS1411 MP surfaces after adding 10 U/mL RNase H and varying RNA/DNA duplex concentrations. Scale bars = 20 µm.
